# Single cell RNA sequencing reveals the landscape of early female germ cell development

**DOI:** 10.1101/2020.05.09.085845

**Authors:** Zheng-Hui Zhao, Jun-Yu Ma, Tie-Gang Meng, Zhen-Bo Wang, Wei Yue, Qian Zhou, Sen Li, Xie Feng, Yi Hou, Heide Schatten, Xiang-Hong Ou, Qing-Yuan Sun

## Abstract

Female germ cell development consists of complex events including sex determination, meiosis initiation, retardation and resumption. During early oogenesis, the asynchrony of the transition from mitosis to meiosis results in heterogeneity in the female germ cell populations at a certain embryonic stage, which limits the studies of meiosis initiation and progression at a higher resolution level. Here, we investigated the transcriptional profiles of 19363 single germ cells collected from E12.5, E14.5 and E16.5 mouse fetal ovaries. Clustering analysis identified seven groups and defined dozens of corresponding transcription factors, providing a global view of cellular differentiation from primordial germ cells towards meiocytes. Further, we explored the dynamics of gene expression within the developmental trajectory with special focus on the mechanisms underlying meiotic initiation. We found that *Dpy30* may be involved in the regulation of meiosis initiation at the epigenetic level. Our data provide key insights into the transcriptome features of peri-meiotic female germ cells, which offers new information not only on meiosis initiation and progression but also on screening pathogenic mutations in meiosis-associated diseases.

---

Primordial germ cells (PGCs) development undergo specification, migration and differentiation processes to generate the gametes. The specification of PGCs is initiated at E6.25 in mice, and they are easily identified at E7.5 by alkaline phosphatase staining (1). Shortly after specification, the PGCs begin migration from the posterior primitive streak to the genital ridges via the hindgut (2). After arriving at the genital ridges at around E10.5, the female PGCs proliferate with an incomplete cytoplasmic mitosis to form oogonia cysts (3) and initiate meiosis asynchronously to become meiocytes at around E12.5 (4). Following this event, the majority of oogonia enter meiosis at around E14.5 and nearly all the female germ cells enter meiosis at around E16.5. Meiosis is initiated with the complex stages of leptotene, zygotene and pachytene, during which germ cells undergo dramatic transitions, such as chromosome repackaging, synapsis and recombination (5). At around E17.5, the primary oocytes begin to arrest at the diplotene stage until LH induces fully-grown oocytes to resume meiosis I after puberty (6). During the prolonged oogenesis, only a small fraction of oocytes ultimately mature and the majority of immature oocytes enter programmed apoptosis (7).

The female germ cell development is guided by stage-specific gene expression patterns, which are primarily controlled by transcription factors (TFs) (8). Dynamic transcriptional patterns shape the unique transcriptional environment in regulatory elements (8) that could further recruit stage-specific TFs to instruct early oogenesis. Although several groups have partially mapped the transcriptional profiles of fetal female germ cells through bulk RNA-sequencing (9-12), the underlying TFs that participate in the early oogenesis have not been fully explored and identified due to evident heterogeneity of female germ cell populations. In addition, although meiosis initiation depends on the stimulation of retinoic acid (RA) that could further activate the meiotic initiator STRA8 (13), the regulatory network between RA and STRA8 remains unclear.

To explore the regulatory mechanisms in early oogenesis at a single cell level, we performed high throughput and unbiased single-cell RNA sequencing on female germ cells at E12.5, E14.5 and E16.5, which respectively correspond to the early meiotic stage (numerous mitotic cells and a few meiotic cells), middle meiotic stage (a few mitotic cells and numerous meiotic cells) and late meiotic stage (nearly all the cells are meiocytes) in early oogenesis. We sequenced 19363 female germ cells and grouped them into 7 clusters. All the clusters were defined according to the known marker genes (14) and developmental trajectory. Moreover, the relevant TFs important for early oogenesis and the transition from mitosis to meiosis were identified. Finally, *Dpy30*, an epigenetic factor, may bridge the RA and STRA8 to regulate meiosis initiation.

## MATERIALS AND METHODS

### Ethics statement

The animal handling procedures were approved by the Animal Care and Use Committee of the Institute of Zoology at the Chinese Academy of Sciences.

### Sample collection

Pou5f1-eGFP-positive germ cells were isolated from E12.5, E14.5 and E16.5 female gonads (Pou5f1-eGFP JAXR mice stock number 004654 X WT C57BL/6J). For each stage, the gonads were microscopically dissected and pooled from 20-30 female embryos. The pooled gonads were treated with 500 µl of Accutase Cell Detachment Solution (Millipore #SCR005) for 10-20 min at 37 °C, and the digestion reaction was quenched after adding 500 µl L15 medium (Gibco # 21083027) with 10% FBS. The cells were filtered through 30 µm Pre-Separation Filters (MiltenyiBiotec #130-041-407) to obtain single cell suspension for fluorescence-activated cell sorting (MoFLo XDP, Beckman Coulter). The mouse female germ cells at each time point were collected after MoFLo XDP cell sorting as eGFP-positive fraction.

### Single-cell library preparation and sequencing

Libraries were prepared using 10X Genomics platform and processed following the manufacturer’s specifications. Briefly, each cell and each transcript are uniquely barcoded using a unique molecular identifier (UMI) and cDNA ready for sequencing on Illumina platforms is generated using the Single Cell 3’ Reagent Kits v2 (10X Genomics). Libraries were sequenced on an Illumina NovaSeq 6000 (Illumina, San Diego) with a read length of 26 bp for read 1 (cell barcode and UMI), 8 bp i7 index read (sample barcode), and 98 bp for read 2 (actual RNA read). Reads were first sequenced in the rapid run mode, allowing for fine-tuning of sample ratios in the following high-output run. Combining the data from both flow cells yielded approximately > 40,000 reads per single-cell.

### Data processing and analysis

Fastq data generated by Illumina NovaSeq 6000 sequencing were processed and mapped to the mouse genome (mm10) with the Cell Ranger software (v2.1.0) with default parameters. Then the Cell Ranger analysis results were used as input for the R package Seurat (15). To check the quality of the single cell data and to remove any multiplets, we performed Seurat-based filtering of cells based on three criteria: number of detected genes per cell, number of UMIs expressed per cell and mitochondrial content, using the following threshold parameters: nFeature_RNA (1500 to 5500), nCount_RNA (between -inf and 40,000), and percent.mt (< 5%). Normalization was performed as described in the package manual (https://satijalab.org/seurat/v3.1/) (16). Next, we combined and scaled the scRNA-seq data of female germ cells, clustered the cells into clusters using Louvain algorithm (17), and visualized the cell clusters with UMAP. The differentially expressed genes were determined by the FindMarkers function. Gene set enrichment analysis was performed with clusterProfiler R package (18).

### Cell trajectory analysis

Single-cell pseudotime trajectories were constructed with the Monocle 2 package (v2.8.0) (19) according to the provided documentation (http://cole-trapnell-lab.github.io/monocle-release/). Ordering genes were identified as having high dispersion across cells. The discriminative dimensionality reduction with trees (DDRTree) method was used to reduce data to two dimensions. Gene sets identified from the destiny analysis were clustered and visualized using the plot_genes_in_pseudotime function.

### Antibodies

Mouse monoclonal anti-DDX4 antibody used for immunofluorescence was purchased from Abcam (ab27591); Rabbit polyclonal anti-DPY30 antibody used for immunofluorescence was purchased from Abcam (ab126352); Rabbit polyclonal anti-POU5F1 antibody used for immunofluorescence was purchased from Santa Cruz (sc-9081); Rabbit polyclonal anti-STRA8 antibody used for immunofluorescence was purchased from Abcam (ab49405); Alexa Fluor 488-conjugated antibody and Alexa Fluor 594-conjugated antibody were purchased from Life Technologies.

### Immunofluorescence

Immunofluorescence was performed as previous described with small modifications (20). For whole-mount staining, dissected gonads were fixed for 20 min in 4% paraformaldehyde in PBS (pH 7.4) and permeabilized with 1% Triton X-100 in PBS for 30 min on ice. Fixed embryos were blocked overnight at 4 □ in PBS containing 0.1% Triton X-100, 10% BSA and 5% normal donkey serum, and were then incubated with primary antibodies in blocking solution overnight at 4 □. Gonads were washed three times for 1 h in PBS containing 0.1% Triton X-100 and 2% BSA before application of secondary antibodies. For detection, secondary antibodies were diluted 1:500 in blocking solution and gonads were incubated overnight at 4 □ followed by three washing steps for 1 h in PBS with 0.1% Triton X-100. Finally, gonads were stained briefly with DAPI and mounted in Vector.

### Availability of data and materials

The high-throughput sequencing data in this study have been deposited in the Gene Expression Omnibus (GEO) database under accession number GSE130212.

## RESULTS

### Isolation and sequencing of female mouse germ cells

To investigate the regulation of meiosis initiation at a single cell level, we collected female germ cells from gonads of E12.5, E14.5 and E16.5 female Tg (Pou5f1-eGFP) mice. Tg (Pou5f1-eGFP) mice express eGFP under Pou5f1 distal enhancer, and the eGFP-positive germ cells from the gonads were collected using fluorescence-activated cell sorting and individual cells were captured and processed with the 10x Genomics’ Chromium System (Fig. 1A, Fig. S1A). We sequenced the female germ cells in an unbiased manner using the high-throughput single-cell RNA-sequencing method. As a result, we successfully sequenced 5103, 9506, 5910 germ cells from E12.5, E14.5, and E16.5 female fetuses, respectively (Fig. S1B). After filtering out somatic cells and low data quality germ cells, 19363 germ cells remained for subsequent analysis (Fig. S1C).

**Fig. 1.**
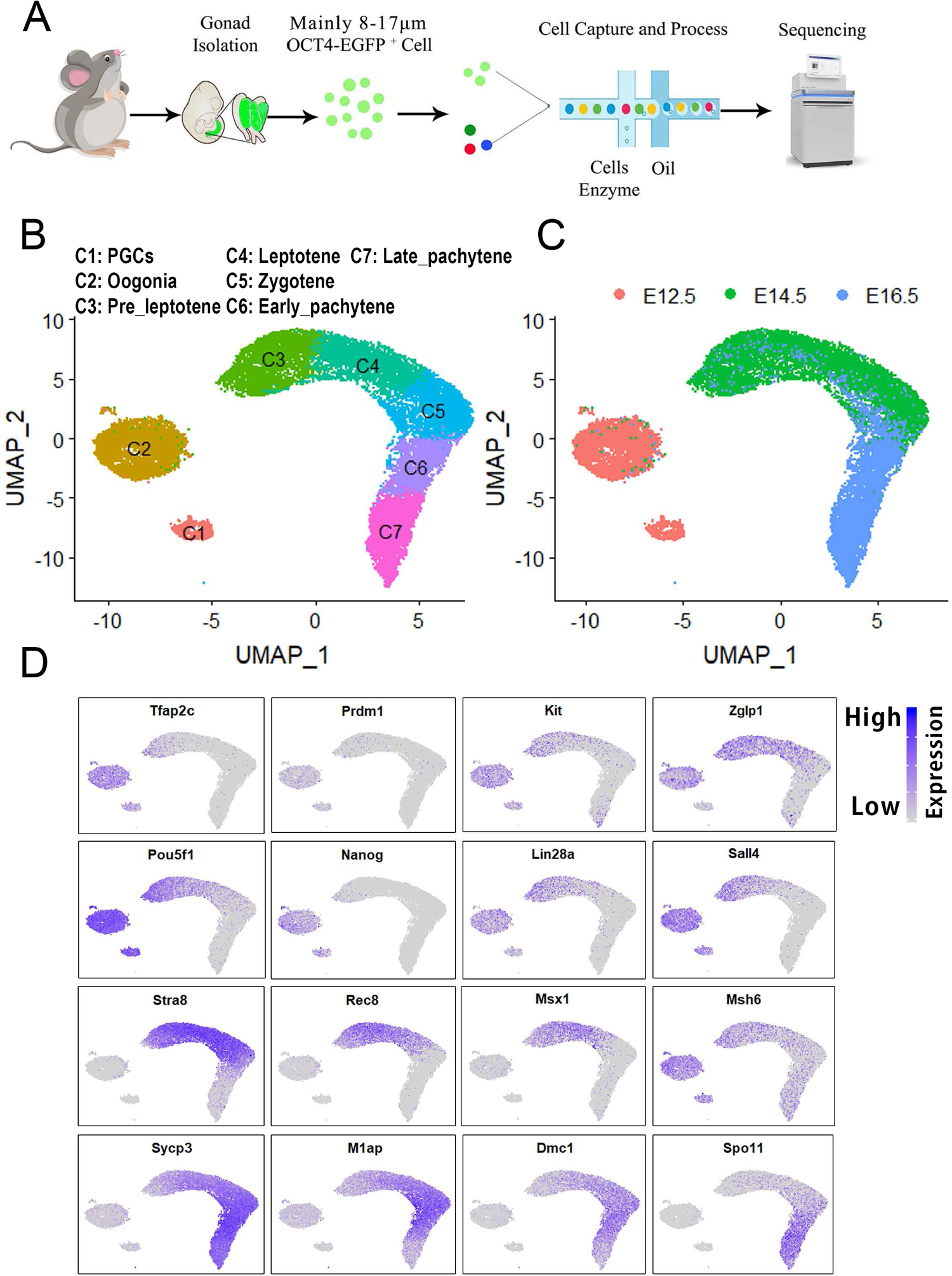
Clustering of the female germ cells during fetal ovarian development. (A) Experimental workflow. XX gonads at E12.5, E14.5 and E16.5 were collected, Pou5f1-eGFP+ cells were sorted by FACS, single-cell captured, and harvested cDNA was processed for libraries, and sequenced. (B) Two-dimensional UMAP representation of the 19363 XX Pou5f1-eGFP+ single cells. Cells were colored according to cluster identity as shown in the Fig. key. (C) UMAP plot of single cell transcriptome data with cells colored by embryonic stages as indicated in the Fig. key. (D) Expression levels of selected markers projected on the UMAP plot. Blue indicates high expression and gray indicates low or no expression, as indicated in the Fig. key.

### Graph clustering identifies seven transcriptional clusters in the female germ cells

To characterize the cell types of female germ cells, we grouped the germ cells into seven clusters (C1-C7) projected on the UMAP plot (Fig. 1B). Each time point of E12.5, E14.5 or E16.5 contains several cell clusters, indicating the evident heterogeneity and developmental asynchrony in female germ cell populations (Fig. 1C). Cluster identity was defined based on known cell-type markers and differentially expressed genes (Fig. 1D, Fig. S1D). The cluster 1 represents primordial germ cell populations, as most cells in this cluster express early PGC markers such as *Prdm1, Tfap2c*, and *Kit* (21), while late germ cell-specific genes such as *Dazl* and *Ddx4* are still lowly expressed (Fig. 1D, Fig. S1E). Also, the cells in C1 express some pluripotent genes such as *Pou5f1, Nanog, Sall4* and *Lin28a*. Moreover, *Lefty1* and *Lefty2* were specifically expressed in C1 cluster, which can be used as new markers for mouse PGCs (Fig. S1E). The oogonia (C2) share the expression of several pluripotent genes with PGCs, such as *Pou5f1* and *Nanog*, but begin to significantly express *Zglp1* that is sufficient to activate the oogenic programs (Fig. 1D) (22), indicating that PGCs and oogonia coexist in E12.5 gonads. The C3 was likely to be pre-leptotene cells, as the cells began to express the meiotic genes, including *Stra8, Rec8, Msx1* and *Msh6*. Additionally, the cells in C3 also expressed certain pluripotent genes such as *Lin28a* and *Sall4* (Fig. 1D). The C4-C7 corresponded to leptotene, zygotene, early_pachytene and late_pachytene cells, respectively, as these cells expressed meiosis markers, such as *Sycp3, M1ap, Spo11* and *Dmc1*, respectively (14).

Furthermore, several stage specific master transcription factors such as *Sox2, Sox4, Dmrt1* and *Dmrtc2* may play important roles in mitosis to meiosis transition and female germ cell development (Fig. S1E). Collectively, with our single-cell RNA sequencing experiments on female germ cells, 7 clusters were identified and they exhibited distinct gene expression patterns.

### Pseudotime trajectory identifies the dynamics of gene expression across female germ cell populations

To further explore the early female germ cell development, we performed monocle trajectory analysis on female germ cells and ordered them along a pseudotime. Notably, the clusters (C1-C7) showed a continuous trajectory, which recapitulated the temporal order of early oogenesis. The PGCs (C1) clustered on one end of the pseudotime axis (the left side), directly following the mitotic oogonia and different stages of meiocytes (Fig. 2A). Additionally, the overlap in cell clusters and embryonic stages indicated that the transcriptome changed continuously in germ cell development (Fig. 2A,B). The reconstruction of cell lineages allowed us to define the transition states from mitosis to meiosis and meiosis progression. By ordering the cells along a pseudotime, we identified several typical genes that expressed at certain stages during female germ cell development. For example, the expression level of meiosis-associated genes increases after the downregulation of pluripotent genes such as *Nanog* (Fig. 2C). And the *Rec8* exhibited the peak expression in pre-leptotene cluster, indicating that *Rec8* may be involved in mitosis to meiosis transition. Furthermore, hierarchical clustering analysis of the ordering genes (rows) while organizing the germ cells (columns) along the pseudotime trajectory revealed 4 distinct gene cohorts. Gene ontology analysis of these grouped genes identified multiple biological processes, which are consistent with the early female germ cell development (Fig. 2D). As expected, “anterior/posterior pattern specification”, “mitosis” and “meiosis” were significantly-enriched gene ontology (GO) categories during female germ cell (FGC) development. With this analysis, we showed that the mitosis to meiosis transition of the female germ cells displayed dramatic changes by downregulating the pluripotent genes and rapidly upregulating meiocyte-specific genes.

**Fig. 2.**
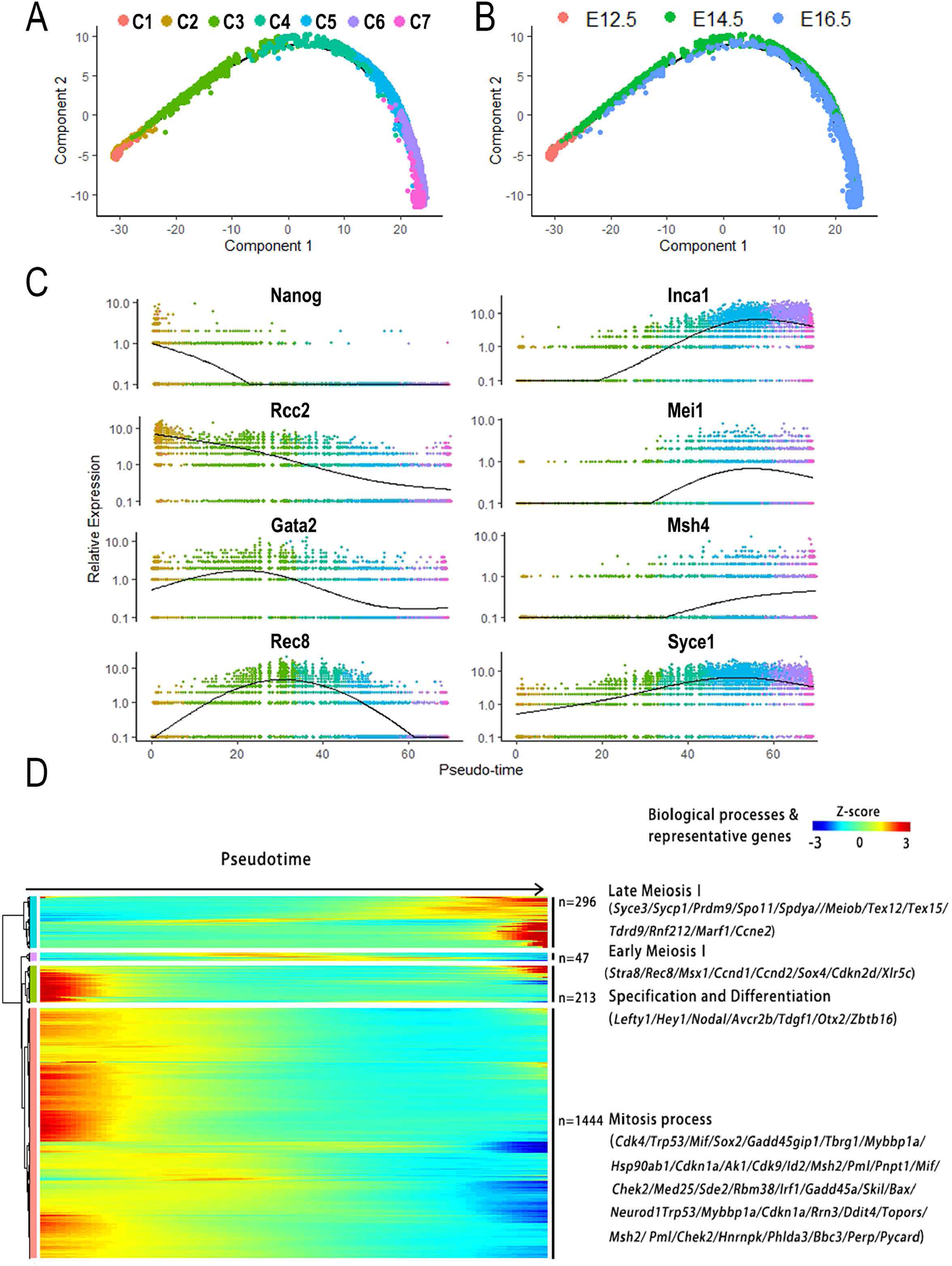
Female germ cell lineage reconstruction and gene expression. (A) Pseudotime trajectories of the female germ cells as indicated in the Fig. key. (B) Pseudotime trajectories of the female germ cells colored by embryonic stages as shown in the Fig. key. (C) Monocle pseudotime trajectory expression pattern of representative genes during transition from mitosis to meiosis. x-axis represents pseudotime, and y-axis represents relative expression levels. (D) Hierarchical clustering of ordering genes exhibiting differential expression (n = 2000) across female germ cell populations.

### Dynamic transcriptional factors instruct early oogenesis

To demonstrate transcriptome changes during early oogenesis, we selected 3575 differentially expressed genes from seven clusters using FindMarkers function in Seurat R package (16). And these differentially expressed genes changed strikingly during mitosis to meiosis transition, but gradually during meiosis progression (Fig. S1D, Fig. S2A-C). Among these genes, we selected 27 highly variable transcription factors that exhibited stage-specific expression patterns and are generally clustered into three groups (Fig. 3A). The first group TFs, such as *Rest* and *Trp53*, mainly expressed in PGCs and oogonia clusters, which may play essential roles in the germ cell proliferation. The second group TFs like *Msx1, Msx2, Gata2, Cdx2, Sox4* and *Bmyc* showed expression peaks specifically in the cluster of pre-leptotene stages, indicating that these TFs may be involved in meiosis initiation. The third group TFs, like meiotic gene *Dmrtc2*, mainly expressed in meiocytes, which may play an important role in meiosis progression. *Taf4b* had a similar expression pattern as *Dmrtc2*, which is essential for female germ cell development and *Taf4b* null females are sterile, indicating that it is required for oocyte development (23, 24).

**Fig. 3.**
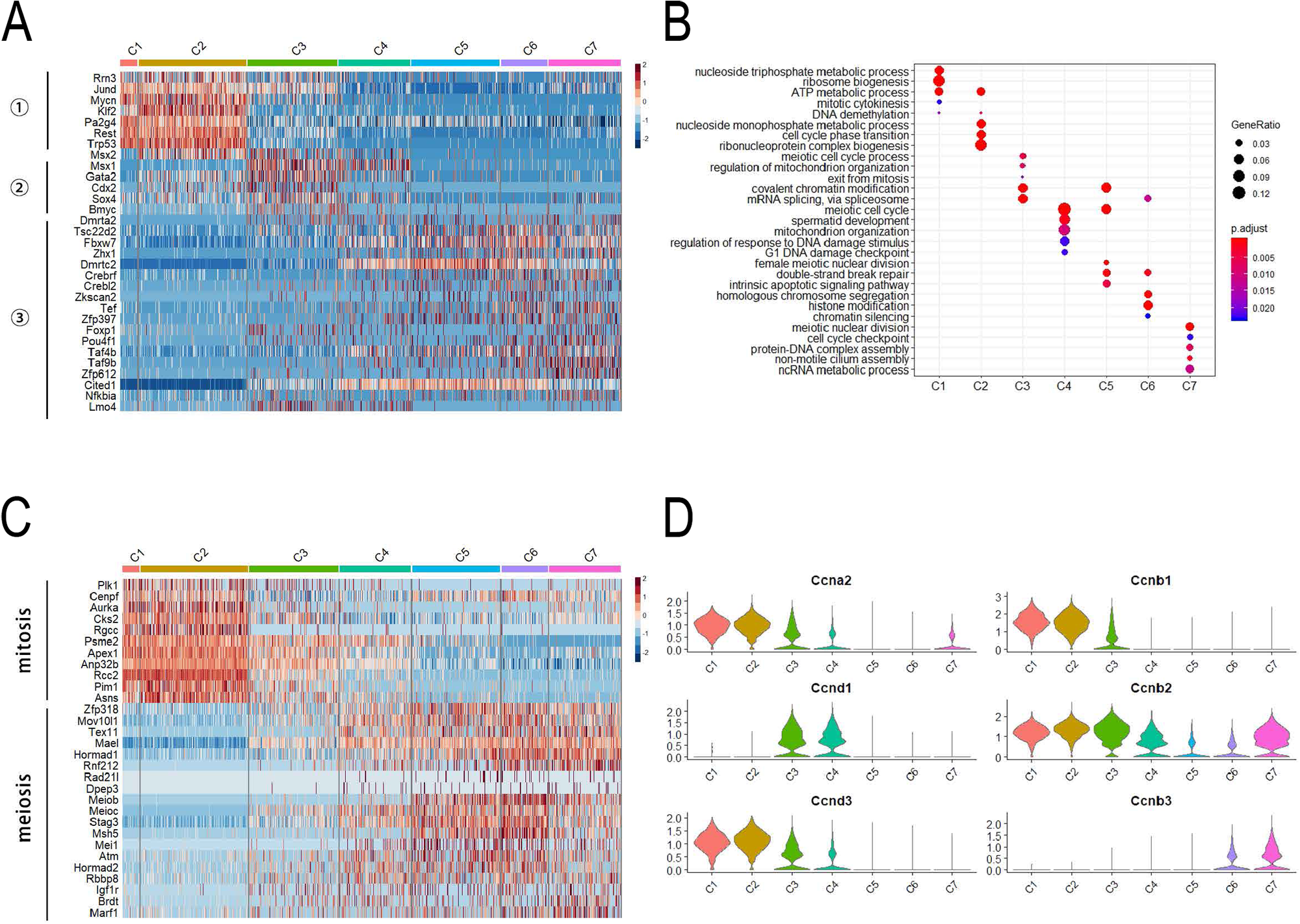
The regulation of mitosis to meiosis transition in female germ cells. (A) Heatmap of highly variable transcription factor genes during germ cell development. (B) Top enriched biological processes of the differentially genes from each of the seven cell clusters. (C) The expression patterns of mitotic and meiotic cell cycle genes. (D) Violin plot of the differentially expressed cyclin genes.

To further dissect the differences in gene expression profiles, we identified 660, 856, 183, 138, 561, 609, and 568 differentially expressed genes among these seven clusters, respectively, and performed gene ontology analysis of these genes. GO categories indicated that the C1 and C2 were mitotic cells (“ribosome biogenesis” and “mitotic cytokinesis”), whereas the C3-C7 were meiocytes that express genes related to meiotic recombination and DNA damage repair (‘‘meiotic cell cycle’’, ‘‘double-strand break repair’’ and “homologous chromosome segregation”) (Fig. 3B). Interestingly, we found that some meiocytes expressed genes such as *Rec8, Dmrtc2* and *Sycp3* also related to spermatid development processes, indicating that the male and female gametes likely share certain mechanisms in meiosis progression (Fig. 3B). Additionally, dynamic cell cycle regulation is crucial for mitosis to meiosis transition. To further explore the mechanisms of the meiosis initiation, we selected highly variable mitotic and meiotic cell cycle genes from all clusters. Consistent with the above cluster identification, mitotic cell cycle genes were mainly expressed in PGCs and oogonia clusters, whereas meiotic cell cycle genes were expressed in meiocytes (C3-C7) (Fig. 3C). In addition, cyclin genes showed different expression patterns. *Ccna2, Ccnb1/2* and *Ccnd3* were highly expressed between C1 and C3 clusters and subsequently decreased during the progression of meiosis, which suggests that their functions are restricted to the early mitosis phase (Fig. 3D). Moreover, the down-regulation of *Ccnb1/2* in early meiosis prophase may be essential for meiosis□ progression. The expression level of *Ccnd1* was low in mitotic PGCs and oogonia, but sharply increased in early meiotic cells, and then decreased during late meiosis phase, whereas *Ccnb3* began to be expressed from the pachytene stages on (Fig. 3D). In addition, the cyclin-associated genes also exhibited different expression patterns. The *Cdk1* and *Cdk4* mainly expressed in C1-C3, which may be involved in the regulation of cell proliferation. In addition, *Cdkn2a* (p16) and *Cdkn2d* (p19), the inhibitors of *Cdk4*, increased consistently with *Ccnd1*, which indicates that *Cdkn2a* and *Cdkn2d* may play a role in balancing the functions of *Ccnd1* in the early meiosis process (Fig. S2D). Moreover, the *Cdkn1a* (p21), whose expression decreased during the transition from mitosis to meiosis, was consistent with the down-regulation of pluripotency markers (Fig. S2E), reflecting the trajectory of oogenesis. Collectively, these data indicate that a dynamic transcriptome regulates early oogenesis in a stage-specific manner.

### Dynamics of epigenetic factors during female germ cell development

To explore the functions of epigenetic factors during early oogenesis, we analyzed the expression patterns of epigenetic factors that participate in DNA methylation and histone modifications in the seven clusters. As oogenesis continues, the level of DNA methylation decreases and the lowest level is maintained until birth (25). During early oogenesis, the *Tet1* may be the only erasure for DNA demethylation, with minimally expressed de novo DNA methylation enzymes and the other two erasures, *Tet2* and *Tet3* (Fig. 4A, Fig. S3A). In addition, the relative high expression of *Dnmt1* and *Uhrf1* in female germ cells may likely be involved in the maintenance of DNA hypomethylation status in early oogenesis stages (Fig. 4A). In contrast to early embryogenesis, the DNA demethylation in oogenesis also involves imprinted regions. Therefore, the maternal and paternal imprinted genes exhibited dynamic expression patterns, and the differentially expressed imprinted genes are listed in Fig. S3B. The expression level of *Peg3* was high in mitotic PGCs and oogonia, which may be related to the regulation of cell proliferation, while other maternal imprinted genes, such as *Kcnq1ot1*, mainly were expressed in meiocytes, suggesting their important roles in the regulation of meiosis progression (Fig. 4B). On the other hand, the paternal imprinted genes, such as *Phlda2* and *Gnas*, also showed unique expression patterns during early oogenesis. The *Phlda2* was specifically expressed in oogonia and pre-leptotene cells, which may be involved in the regulation of mitosis to meiosis transition. In addition, *Gnas* was highly expressed in meiocytes, indicating that it may participate in the regulation of the meiosis process (Fig. 4B).

**Fig. 4.**
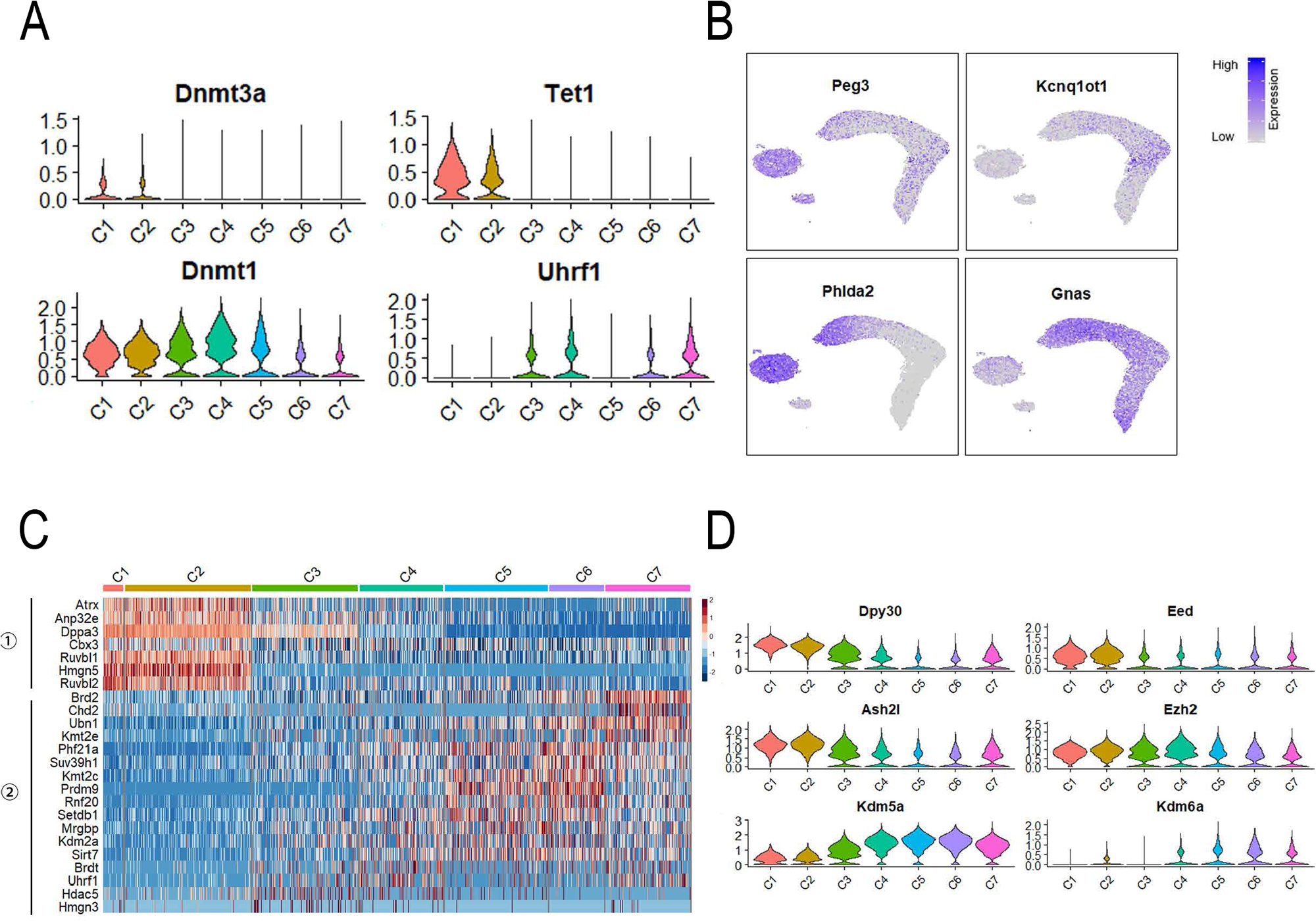
The epigenetic regulation in female germ cell development. (A) The expression patterns of DNA methylation modifiers during germ cell development. (B) The differentially expressed maternal and paternal imprinting genes. (C) Heatmap of highly variable covalent chromatin modification-associated genes during germ cell development. (D) The expression patterns of H3K4me3 and H3K27me3 associated modifiers.

As previously reported, histone modifications play important roles in mitosis to meiosis transition (10, 26). To explore the functions of histone modifications in early oogenesis, we selected differentially expressed covalent chromatin modification-related genes that were clustered into two groups (Fig. 4C). Interestingly, the first group of genes were mainly expressed in mitotic PGCs and oogonia clusters, whereas the second group of genes were highly expressed in meiocytes, which suggests that the covalent chromatin modifications, such as H3K4me3 and H3K27me3, undergo a process of redistribution during the transition from mitosis to meiosis (Fig. 4C) (10). To further support this idea, we also determined the expression patterns of writers and erasures for the H3K4me3 and H3K27me3 (Fig. 4D). The *Dpy30* and *Ash2l* are the subunits of MLL histone methyltransferase complexes that catalyze H3K4me3 directly. The expression levels of *Kdm5a* increased gradually after entering meiosis. Moreover, the PRC2 subunits *Eed* and *Ezh2*, which catalyze H3K27me3, maintained stable expression levels during early oogenesis. However, the H3K27me3 de-methyltransferase *Kdm6a* exhibited a higher expression level in zygotene and pachytene stages (Fig. 4D). The dynamic expression patterns of these enzymes may suggest the redistribution of H3K4me3 and H3K27me3. On the other hand, although the level of DNA methylation was low in early oogenesis, there was not an obvious increase in the gene expression level, which suggests that other mechanisms may be responsible for chromatin silencing. The *Ehmt2, Setdb1* and *Suv39h1* presented a higher expression level in pre-leptotene, zygotene, and pachytene stages, respectively (Fig. S3C), which catalyze H3K9me2 and H3K9me3, and both of the modifications may compensate for the low level of DNA methylation.

### Dpy30 may be responsible for meiosis initiation

Meiosis initiation is a critical event during female germ cell development. Recent study has shown that BMP-ZGLP1 pathway could specify the oogenic program, and RA pathway contribute to the overall maturation of the oogenic program as well as to the repression of the PGC program (22). To investigate the mechanisms of meiosis initiation, we performed reclustering of the oogonia cluster with high resolution and four clusters (Oogonia 1-4) were identified (Fig. 5A). The majority of oogonia cells derived from E12.5 ovaries, whilst E14.5 and E16.5 ovaries also contain a few oogonia cells (Fig. 5B). Furthermore, cluster identity was defined based on differentially expressed genes (Fig. 5C). Although *Utf1* showed a uniform expression in oogonia cells, the pluripotent gene *Dppa5a* was highly expressed in the middle region of the oogonia cluster. In contrast to *Dppa5a, Dazl* exhibited higher expression level in the peripheral region of the oogonia cluster (Fig. 5D). Additionally, the cells in oogonia 4 cluster highly expressed late PGC marker gene *Dazl* and meiotic marker gene *Stra8*, suggesting that the cells in oogonia 4 begin to entry meiosis (Fig. 5C-D).

**Fig. 5.**
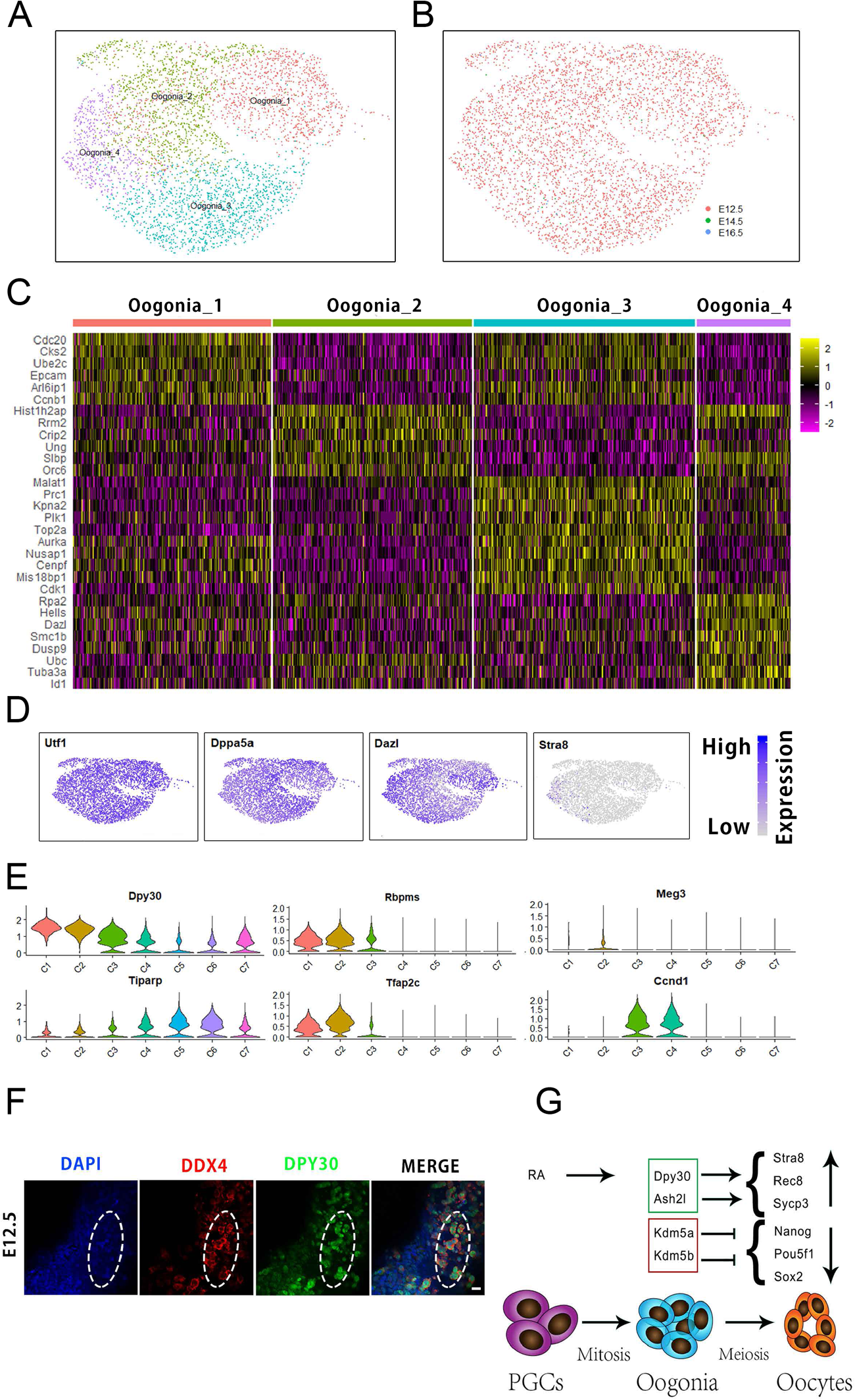
The regulation of meiosis initiation. (A) Reclustering of the oogonia cluster (C2) at a higher resolution. (B) UMAP plot of oogonia transcriptome data with cells colored by embryonic stages. (C) Heatmap of differentially expressed genes across four oogonia subpopulations. (D) The expression patterns of pluripotency and meiosis-associated genes. (E) The expression patterns of retinoic acid target genes in female germ cells. (F) Immunofluorescence staining of DDX4 and DPY30 in E12.5 female gonads. Scale bars, 20 μm. (G) The predicted model for the regulation of meiosis initiation.

To further explore the mechanisms of RA pathway in the maturation of the oogenic program and repression of the PGC program, we selected several RA target genes that were differentially expressed in early oogenesis (Fig. 5E). Previous study has indicated that *Dpy30* is crucial for normal H3K4me3 increase at the loci of developmental genes for RA-mediated cell fate transition (27). In this study, *Dpy30* was highly expressed in oogonia cluster, which may catalyze the H3K4me3 in the promoters of meiotic genes, including *Stra8, Rec8* and *Sycp3* (10). Also, immunofluorescence analysis revealed that DPY30 exhibited robust expression in E12.5 female germ cells (Fig. 5F). Furthermore, *Ash2l* exhibited similar expression pattern to *Dpy30* (Fig. 4D), which was also involved in the establishment of H3K4me3, suggesting similar roles to *Dpy30*. Meanwhile, the expression of Kdm5a in the oogonia stage may be involved in the erasure of H3K4me3 in the promoters of pluripotent genes, including *Pou5f1* and *Nanog* (Fig. 4D). Collectively, these data indicated that *Dpy30, Ash2l* and *Kdm5a* may switch on the meiotic genes and switch off the pluripotent genes to promote meiosis initiation and repress PGC program at the epigenetic level (Fig. 5G).

## DISCUSSION

Female germ cell development is a complex process involving the transition from mitosis to meiosis and meiosis progression. In this study, we aimed to shed light on the mechanisms regulating these processes through high throughput single cell RNA sequencing of female germ cells. We obtained single cell transcriptome data and identified 3 distinct germ cell stages that are PGCs, mitotic oogonia and meiotic oocytes, respectively. Moreover, meiotic oocytes also contain several subsets that form pre-leptotene to late-pachytene clusters. Our transcriptome data provide new insights into the molecular signatures associated with the assignment of each cell cluster. Understanding the transition from mitosis to meiosis in early oogenesis is essential for the application of induced gametes.

Here, we reconstructed the dynamic and temporal trajectory of female germ cell development using monocle2 R package (19), which is consistent with the identified cell clusters. With this analysis, we found that early oogenesis presents a linear developmental process by downregulating the pluripotent genes and upregulating meiotic genes. Although two distinct clusters were computationally obtained (Clusters PGCs and Oogonia), an alternative interpretation was that these two populations were similar, and simply expressed several different genes including *Lefty1, Lefty2* and *Phlda2* (Fig. 4B, Fig. S1E). In addition, dozens of differentially expressed TFs were detected during early oogenesis. Of these, several TFs belonged to the homeobox family containing helix-turn-helix motifs, which could directly bind to the regulatory elements of target genes and promote transcription. For example, *Cdx2* that is required for the differentiation of trophectoderm and is inhibited by *Pou5f1* in the inner cell mass at the blastocyst stage (28, 29), is exclusively expressed in pre-leptotene cluster, suggesting its roles in the regulation of meiosis initiation. Furthermore, *Cyp26a1*, the direct targeted gene of *Cdx2*, could affect the retinoid signaling in the developing embryo, indicating that it is closely related to meiosis initiation (30). Moreover, various TF cofactors including *Cited1, Nfkbia*, and *Lmo4* were also identified in female germ cells (Fig. 3A), and these cofactors could interact with numerous TFs that are highly expressed in female germ cells to regulate early oogenesis (31-33).

Dynamic cell cycle regulation is crucial for female germ cell development. Our data indicated that the cell cycle genes in meiocytes underwent complex regulations and they were very different compared to those in somatic cells. In somatic cells, *Ccnb1* is essential for chromatin condensation and mitosis initiation, and it is degraded by the anaphase promoting complex during the metaphase/anaphase transition (34). In our data we found that the *Ccnb1* mRNA was downregulated from the pre-leptotene stage, which suggests that *Ccnb1* is dispensable for the meiocytes’ chromatin remodeling. In addition, the *Ccnd1* targeted by RA is highly expressed in the pre-leptotene cluster, which suggests that it may be involved in the regulation of meiosis initiation.

Genome-wide DNA demethylation takes place during PGC development, and nearly all the genomic elements appear to be uniformly hypomethylated (35). In contrast to early embryogenesis, the DNA demethylation in PGC development also erases the DNA methylation in imprinting control regions (ICRs), and the unmethylated state is maintained until birth during early oogenesis. Therefore, the maternal and paternal imprinted genes are co-expressed during early oogenesis. Although there is no direct evidence showing a relationship between imprinted genes and the regulation of oogenesis, the differentially expressed imprinted genes, such as *Peg3, Kcnq10t1* and *Gpr1*, may be related to regulate the appropriate stage of female germ cells (36-38). On the other hand, there is no obvious expression of development-associated genes in the global demethylation environment, suggesting that repressed histone modifications may be responsible for chromatin silencing instead. During early oogenesis, the PRC2 complex-associated genes and *Ehmt2, Setdb1* and *Suv39h1* were expressed in a stage-specific manner, which may play an essential role in maintaining chromatin silencing.

Meiosis initiation is a critical event during early oogenesis; however, detailed mechanisms that regulate the transition from mitosis to meiosis remain unclear. In this study, we found that several critical TFs, including *Msx1, Msx2, Gata2, Cdx2, Bmyc* and *Sox4*, specifically were expressed at the pre-leptotene stage (Fig. S3D), suggesting their possible roles in meiosis initiation. However, *Msx1* knockout had no impact on meiosis initiation, but double knockout of *Msx1* and *Msx2* could result in decreased *Stra8* expression and the number of *Sycp3* positive meiocytes, which reveals a compensatory mechanism between *Msx1* and *Msx2* in regulating *Stra8* expression (39). Moreover, *Gata2* has a similar expression pattern as *Msx2*, and it is generally consistent with its protein expression, suggesting its role in promoting meiosis initiation (40). Additionally, we detected robust signals of POU5F1 and STRA8 at the E12.5 and E14.5 female gonads respectively, whilst we didn’t detect the signals of STRA8 and POU5F1 at the E12.5 and E14.5 female gonads respectively through whole-mount staining (Fig. S3E). These results indicated that the transition from mitosis to meiosis mainly occurs between these two stages. Furthermore, *Sox4* is also highly expressed at the pre-leptotene stage. Although *Sox4* is essential for male germ cell differentiation, the female fetal germ cells appear unperturbed in *Sox4* knockout mice (41). In addition, H3K4me3 ChIP-seq on E12.5 and E14.5 female germ cells has demonstrated that the level of H3K4me3 at the promoters of meiotic genes increases significantly, whereas that at the promoters of pluripotent genes decreases during the transition from mitosis to meiosis (10). In this study, we found that *Dpy30*, targeted by RA, is likely to catalyze H3K4me3 at the promoters of meiotic genes such as *Stra8, Rec8* and *Sycp3*. Meanwhile, *Kdm5a* may be involved in the erasure of H3K4me3 at the promoters of pluripotent genes such as *Pou5f1*, and *Nanog*. All of these epigenetic factors may play important roles in switching on meiotic genes and switching off pluripotent genes to further promote meiosis initiation and repress PGC program.

In conclusion, using the scRNA-seq method, we have obtained the expression profiles of 19363 female germ cells from PGCs to oogonia and then to meiotic pachytene oocytes. Data presented in this study provide us with a comprehensive view to understand the regulatory network for meiosis initiation and progression. In addition, we listed the meiosis-related genes from the perspectives of the cell cycle, transcription factors and DNA damage repair-associated genes. These findings will expand our knowledge of female germ cell meiosis and might also be helpful for screening genes that may be applied to in vitro culture of gametes.

## ACKNOWLEDGMENTS

We thank Dr. Jin-Song Li from Shanghai Institute of Biochemistry and Cell Biology, Chinese Academy of Sciences & University of Chinese Academy of Sciences for providing Pou5f1-eGFP transgene mice. We thank Wen-Bo Liu and Wen-Long Lei from State Key Laboratory of Stem Cell and Reproductive Biology, Institute of Zoology, Chinese Academy of Sciences for their assistance with gonads collection. We thank Fei-Yang Wang from State Key Laboratory of Stem Cell and Reproductive Biology, Institute of Zoology, Chinese Academy of Sciences for their guidance with bioinformatic analysis. We thank Jian Li from State Key Laboratory of Stem Cell and Reproductive Biology, Institute of Zoology, Chinese Academy of Sciences for the comments on the preparation of the manuscript. The authors declare that they have no conflict of interests.

## AUTHOR CONTRIBUTIONS

Q. Sun designed and supervised the study. Z. Zhao conducted the study and analyzed the data. J. Ma assisted for the data analysis. T. Meng, Z. Wang, X. Ou, Q. Zhou, and W. Yue provided the technical support. Z. Zhao, H. Schatten and Q. Sun wrote the paper. All authors read and approved the final manuscript.

## ABBREVIATIONS

DDRTree: discriminative dimensionality reduction with trees
FGC: female germ cell
GO: gene ontology
ICRs: imprinting control regions
PGCs: primordial germ cells
RA: retinoic acid
TFs: transcription factors
UMI: unique molecular identifier

## Figure legends

**Supplementary Fig. S1** Cell isolation, sequencing information, quality evaluation and upregulated genes in each cluster. (A) FACS profiles of Tg(Pou5f1-EGFP^+^) ovary show a reduction of the fluorescence intensity from E12.5 to E16.5 (B) Sequencing information including cell number, mean reads per cell and median genes per cell. (C) Quality evaluation on sequencing data including nFeature_RNA, nCount_RNA and percent.mt. (D) Heatmap of differentially expressed genes used to classify cell types for each cluster compared to all other clusters for the 7 germ cell clusters. (E) Feature plot of critical genes closely related to germ cell development.

**Supplementary Fig. S2** The gene expression patterns of the transition from mitosis to meiosis. (A) Venn diagram of the highly variable genes in each cluster. (B) Heatmap of DNA recombination-associated genes during germ cell development. (C) Heatmap of DNA repair-associated genes during germ cell development. (D) The expression levels of cyclin-associated genes. (E) The expression patterns of pluripotent genes in germ cell development.

**Supplementary Fig. S3** The expression patterns of imprinted genes and histone modification-associated enzymes. (A) The expression levels of DNA methylation modifiers. (B) The differentially expressed maternal and paternal imprinting genes. (C) The expression levels of histone modification-associated enzymes. (D) Violin plot of the critical TFs related to meiosis initiation. (E) Immunofluorescence staining of POU5F1 and STRA8 in female gonads at E12.5 and E14.5 respectively. Scale bars, 100 μm.

## Notes

### Competing Interest Statement

The authors have declared no competing interest.

## REFERENCES

1. Saitou, M., Barton, S. C., and Surani, M. A. (2002) A molecular programme for the specification of germ cell fate in mice. Nature 418, 293–300

2. Tam, P. P., and Snow, M. H. (1981) Proliferation and migration of primordial germ cells during compensatory growth in mouse embryos. Journal of embryology and experimental morphology 64, 133–147

3. Pepling, M. E., and Spradling, A. C. (1998) Female mouse germ cells form synchronously dividing cysts. Development (Cambridge, England) 125, 3323–3328

4. Bowles, J., and Koopman, P. (2007) Retinoic acid, meiosis and germ cell fate in mammals. Development (Cambridge, England) 134, 3401–3411

5. Sanchez, F., and Smitz, J. (2012) Molecular control of oogenesis. Biochimica et biophysica acta 1822, 1896–1912

6. Borum, K. (1961) Oogenesis in the mouse. A study of the meiotic prophase. Experimental cell research 24, 495–507

7. Kerr, J. B., Hutt, K. J., Michalak, E. M., Cook, M., Vandenberg, C. J., Liew, S. H., Bouillet, P., Mills, A., Scott, C. L., Findlay, J. K., and Strasser, A. (2012) DNA damage-induced primordial follicle oocyte apoptosis and loss of fertility require TAp63-mediated induction of Puma and Noxa. Molecular cell 48, 343–352

8. Ng, H. H., and Surani, M. A. (2011) The transcriptional and signalling networks of pluripotency. Nature cell biology 13, 490–496

9. Guo, H., Hu, B., Yan, L., Yong, J., Wu, Y., Gao, Y., Guo, F., Hou, Y., Fan, X., Dong, J., Wang, X., Zhu, X., Yan, J., Wei, Y., Jin, H., Zhang, W., Wen, L., Tang, F., and Qiao, J. (2017) DNA methylation and chromatin accessibility profiling of mouse and human fetal germ cells. Cell research 27, 165–183

10. Lesch, B. J., Dokshin, G. A., Young, R. A., McCarrey, J. R., and Page, D. C. (2013) A set of genes critical to development is epigenetically poised in mouse germ cells from fetal stages through completion of meiosis. Proceedings of the National Academy of Sciences of the United States of America 110, 16061–16066

11. Ng, J. H., Kumar, V., Muratani, M., Kraus, P., Yeo, J. C., Yaw, L. P., Xue, K., Lufkin, T., Prabhakar, S., and Ng, H. H. (2013) In vivo epigenomic profiling of germ cells reveals germ cell molecular signatures. Developmental cell 24, 324–333

12. Soh, Y. Q., Junker, J. P., Gill, M. E., Mueller, J. L., van Oudenaarden, A., and Page, D. C. (2015) A Gene Regulatory Program for Meiotic Prophase in the Fetal Ovary. PLoS genetics 11, e1005531

13. Endo, T., Freinkman, E., de Rooij, D. G., and Page, D. C. (2017) Periodic production of retinoic acid by meiotic and somatic cells coordinates four transitions in mouse spermatogenesis. Proceedings of the National Academy of Sciences of the United States of America 114, E10132–e10141

14. Handel, M. A., and Schimenti, J. C. (2010) Genetics of mammalian meiosis: regulation, dynamics and impact on fertility. Nature reviews. Genetics 11, 124–136

15. Butler, A., Hoffman, P., Smibert, P., Papalexi, E., and Satija, R. (2018) Integrating single-cell transcriptomic data across different conditions, technologies, and species. Nature biotechnology 36, 411–420

16. Satija, R., Farrell, J. A., Gennert, D., Schier, A. F., and Regev, A. (2015) Spatial reconstruction of single-cell gene expression data. Nature biotechnology 33, 495–502

17. Blondel, V. D., Guillaume, J.-L., Lambiotte, R., and Lefebvre, E. (2008) Fast unfolding of communities in large networks. Journal of Statistical Mechanics: Theory and Experiment 2008

18. Yu, G., Wang, L. G., Han, Y., and He, Q. Y. (2012) clusterProfiler: an R package for comparing biological themes among gene clusters. Omics: a journal of integrative biology 16, 284–287

19. Qiu, X., Mao, Q., Tang, Y., Wang, L., Chawla, R., Pliner, H. A., and Trapnell, C. (2017) Reversed graph embedding resolves complex single-cell trajectories. Nature methods 14, 979–982

20. Yokobayashi, S., Liang, C. Y., Kohler, H., Nestorov, P., Liu, Z., Vidal, M., van Lohuizen, M., Roloff, T. C., and Peters, A. H. (2013) PRC1 coordinates timing of sexual differentiation of female primordial germ cells. Nature 495, 236–240

21. Wen, L., and Tang, F. (2019) Human Germline Cell Development: from the Perspective of Single-Cell Sequencing. Molecular cell 76, 320–328

22. Nagaoka, S. I., and Nakaki, F. (2020) ZGLP1 is a determinant for the oogenic fate in mice. 367

23. Falender, A. E., Freiman, R. N., Geles, K. G., Lo, K. C., Hwang, K., Lamb, D. J., Morris, P. L., Tjian, R., and Richards, J. S. (2005) Maintenance of spermatogenesis requires TAF4b, a gonad-specific subunit of TFIID. Genes & development 19, 794–803

24. Falender, A. E., Shimada, M., Lo, Y. K., and Richards, J. S. (2005) TAF4b, a TBP associated factor, is required for oocyte development and function. Developmental biology 288, 405–419

25. Smallwood, S. A., and Kelsey, G. (2012) De novo DNA methylation: a germ cell perspective. Trends in genetics: TIG 28, 33–42

26. Sachs, M., Onodera, C., Blaschke, K., Ebata, K. T., Song, J. S., and Ramalho-Santos, M. (2013) Bivalent chromatin marks developmental regulatory genes in the mouse embryonic germline in vivo. Cell reports 3, 1777–1784

27. Jiang, H., Shukla, A., Wang, X., Chen, W. Y., Bernstein, B. E., and Roeder, R. G. (2011) Role for Dpy-30 in ES cell-fate specification by regulation of H3K4 methylation within bivalent domains. Cell 144, 513–525

28. Niwa, H., Toyooka, Y., Shimosato, D., Strumpf, D., Takahashi, K., Yagi, R., and Rossant, J. (2005) Interaction between Oct3/4 and Cdx2 determines trophectoderm differentiation. Cell 123, 917–929

29. Strumpf, D., Mao, C. A., Yamanaka, Y., Ralston, A., Chawengsaksophak, K., Beck, F., and Rossant, J. (2005) Cdx2 is required for correct cell fate specification and differentiation of trophectoderm in the mouse blastocyst. Development (Cambridge, England) 132, 2093–2102

30. Savory, J. G., Bouchard, N., Pierre, V., Rijli, F. M., De Repentigny, Y., Kothary, R., and Lohnes, D. (2009) Cdx2 regulation of posterior development through non-Hox targets. Development (Cambridge, England) 136, 4099–4110

31. Xu, Y., Luo, X., Fang, Z., Zheng, X., Zeng, Y., Zhu, C., Gu, J., Tang, F., Hu, Y., Hu, G., Jin, Y., and Li, H. (2018) Transcription coactivator Cited1 acts as an inducer of trophoblast-like state from mouse embryonic stem cells through the activation of BMP signaling. Cell death & disease 9, 924

32. Coto, E., Diaz-Corte, C., Tranche, S., Gomez, J., Alonso, B., Iglesias, S., Reguero, J. R., Lopez-Larrea, C., and Coto-Segura, P. (2018) Gene variants in the NF-KB pathway (NFKB1, NFKBIA, NFKBIZ) and their association with type 2 diabetes and impaired renal function. Human immunology 79, 494–498

33. Jin, K., Xiao, D., Andersen, B., and Xiang, M. (2016) Lmo4 and Other LIM domain only factors are necessary and sufficient for multiple retinal cell type development. Developmental neurobiology 76, 900–915

34. Thornton, B. R., and Toczyski, D. P. (2003) Securin and B-cyclin/CDK are the only essential targets of the APC. Nature cell biology 5, 1090–1094

35. Wang, L., Zhang, J., Duan, J., Gao, X., Zhu, W., Lu, X., Yang, L., Zhang, J., Li, G., Ci, W., Li, W., Zhou, Q., Aluru, N., Tang, F., He, C., Huang, X., and Liu, J. (2014) Programming and inheritance of parental DNA methylomes in mammals. Cell 157, 979–991

36. Relaix, F., Wei, X. J., Wu, X., and Sassoon, D. A. (1998) Peg3/Pw1 is an imprinted gene involved in the TNF-NFkappaB signal transduction pathway. Nature genetics 18, 287–291

37. Korostowski, L., Sedlak, N., and Engel, N. (2012) The Kcnq1ot1 long non-coding RNA affects chromatin conformation and expression of Kcnq1, but does not regulate its imprinting in the developing heart. PLoS genetics 8, e1002956

38. Xue, Y., Batlle, M., and Hirsch, J. P. (1998) GPR1 encodes a putative G protein-coupled receptor that associates with the Gpa2p Galpha subunit and functions in a Ras-independent pathway. The EMBO journal 17, 1996–2007

39. Le Bouffant, R., Souquet, B., Duval, N., Duquenne, C., Herve, R., Frydman, N., Robert, B., Habert, R., and Livera, G. (2011) Msx1 and Msx2 promote meiosis initiation. Development (Cambridge, England) 138, 5393–5402

40. Onodera, K., Fujiwara, T., Onishi, Y., Itoh-Nakadai, A., Okitsu, Y., Fukuhara, N., Ishizawa, K., Shimizu, R., Yamamoto, M., and Harigae, H. (2016) GATA2 regulates dendritic cell differentiation. Blood 128, 508–518

41. Zhao, L., Arsenault, M., Ng, E. T., Longmuss, E., Chau, T. C., Hartwig, S., and Koopman, P. (2017) SOX4 regulates gonad morphogenesis and promotes male germ cell differentiation in mice. Developmental biology 423, 46–56

